# Interplay between Two Paralogous Human Silencing Hub (HuSH) Complexes in Regulating LINE-1 Element Silencing

**DOI:** 10.1101/2023.12.28.573526

**Authors:** Zena D. Jensvold, Anna E. Christenson, Julia R. Flood, Peter W. Lewis

**Affiliations:** Department of Biomolecular Chemistry, School of Medicine and Public Health, University of Wisconsin, Madison, WI 53715, USA

## Abstract

The Human Silencing Hub (HuSH) complex is composed of TASOR, MPP8, and PPHLN1 subunits and serves as a conserved protein complex responsible for silencing transposable elements in vertebrate animals. Despite its importance, the regulatory mechanisms and recruitment dynamics governing this complex remain poorly understood. In this study, we have identified a second HuSH complex, termed HuSH2, centered around TASOR2, a paralog of the core TASOR protein in HuSH. Our findings indicate that every subunit in both HuSH and HuSH2 has an important role in achieving precise genomic localization to distinct, non-overlapping genomic loci. We utilized in silico protein structure prediction to simulate the interactions between MPP8 and both TASOR paralogs. Drawing on the insights gained from these predictions, we implemented amino acid substitutions that interfered with the binding of MPP8 to each HuSH complex. Leveraging these MPP8 transgenes and other constructs, we identified an important role played by the relative quantities of HuSH complexes in controlling the activity of LINE-1 elements. Furthermore, our results suggest that dynamic changes in TASOR and TASOR2 expression enable cells to finely tune the extent of HuSH-mediated silencing. Our study provides insights into the intricate interplay between HuSH complexes, illuminating their important role in the regulation of retrotransposon silencing.

**Key Points:**

- The identification of a previously unknown HuSH2 complex, with TASOR2 as its central component.
- HuSH and HuSH2 complexes exhibit unique genomic localization patterns within the human genome.
- Disruption of the delicate balance between the two HuSH complexes results in the desilencing of LINE-1.
- TASOR and TASOR2 engage in a competitive interaction for the HuSH subunit MPP8.
- The localization of MPP8 to either HuSH or HuSH2 sites is intricately regulated by its interaction with TASOR and TASOR2.

## Introduction

Retrotransposable elements constitute a noteworthy component of the vertebrate genomic landscape, as their sequences, derived from retroelements, make up approximately 42% of the human genome. This finding highlights the important role played by reverse transcription products and the reverse transcriptase enzyme in shaping genome composition and regulation. To counter potential genomic instability caused by transposons, cells have evolved intricate silencing mechanisms that involve covalent modifications to histones and DNA, effectively suppressing the activity of these parasitic genetic elements.

The HuSH complex, comprised of TASOR, MPP8, and PPHLN1, assumes a central role in the epigenetic regulation of retrotranspons through association with chromatin and RNA modifying proteins. For example, HuSH associates with SETDB1, facilitating the deposition of H3K9me3, and MORC2, an ATP-dependent chromatin remodeler crucial for transcriptional repression^1,2^. The recruitment of these effectors stimulates the catalysis of histone H3 lysine 9 trimethylation (H3K9me3) and facilitates the formation of heterochromatin^1,3^. The targets of the HuSH complex include endogenous retroviruses (ERVs), LINE-1 (L1) retrotransposons, pseudogenes, and intronless mRNAs transcribed by RNA polymerase II that exceed a length of approximately 1.5 kilobases^1,4,5,6^.

Although a comprehensive understanding of how the HuSH complex is recruited to genomic loci remains incomplete, previous work hints at the complex’s engagement with RNA for effective transposable element repression^6,7^. Additionally, the presence of introns appears to confer a protection against HuSH-mediated silencing of transgenes^6^. Through genome-wide analyses, two categories of HuSH targets have been identified. The first category consists of evolutionarily young transposable elements that exhibit a high abundance of repressive histone modifications^8^. The second category comprises expressed coding and non-coding pseudogenes that lack repressive histone modifications and transposons^6,9^. Although the role of HuSH in silencing is well-established, the specific regulatory mechanisms governing HuSH localization and function are not fully understood.

In this study, we have discovered a variant of the HuSH complex, HuSH2, which contains MPP8 and PPHLN like the canonical complex, but instead is centered around TASOR2. In both HuSH complexes, the stability of the subunits is interconnected, and the loss of any subunit leads to a decrease in protein levels of the other subunits. The abundance of HuSH and HuSH2 complexes is affected by changes in TASOR and TASOR2 levels due to competition between TASOR proteins for a limited amount of MPP8 and PPHLN1. Altering the balance of the two HuSH complexes within cells resulted in the induction of transposable elements such as LINE-1 in human lymphoblast cells. Furthermore, the intricate interplay between the HuSH and HuSH2 complexes involves their specific binding to distinct locations on human chromosomes. Notably, fluctuations in the levels of the core TASOR proteins disrupt the precise localization of these complexes, emphasizing the essential role of this core subunit in maintaining proper genomic organization. By introducing targeted residue substitutions that disrupt the interactions between MPP8 and TASOR or TASOR2, our experimental findings provide compelling evidence that these interactions play a crucial role in governing the localization and silencing mechanisms of HuSH. Our study underscores the important impact of both TASOR proteins on the abundance of the HuSH complex, highlighting their essential role in regulating silencing mechanisms.

## Results

The mechanisms by which the HuSH complex is recruited to genomic regions are still poorly understood, and significant questions remain regarding how this complex effectively silences retrotransposable elements and pseudogenes. Notably, the subunits that constitute the HuSH complex lack apparent enzymatic activity. Instead, the prevailing model proposes that the complex primarily functions as a central hub, enabling important protein-protein interactions necessary for its silencing activities^7^. To identify proteins associated with the HuSH complex, which may contribute to its recruitment and silencing functions, and potentially discover novel core members, we conducted a comprehensive study using a rigorous immunoprecipitation (IP) approach combined with liquid chromatography-tandem mass spectrometry (LC-MS/MS) analysis. Specifically, we used a stably expressed epitope-tagged transgenes of the known HuSH core complex members, namely TASOR, MPP8, and PPHLN1, in HEK293 cells (**Figure 1B-D**).

**Figure 1:**
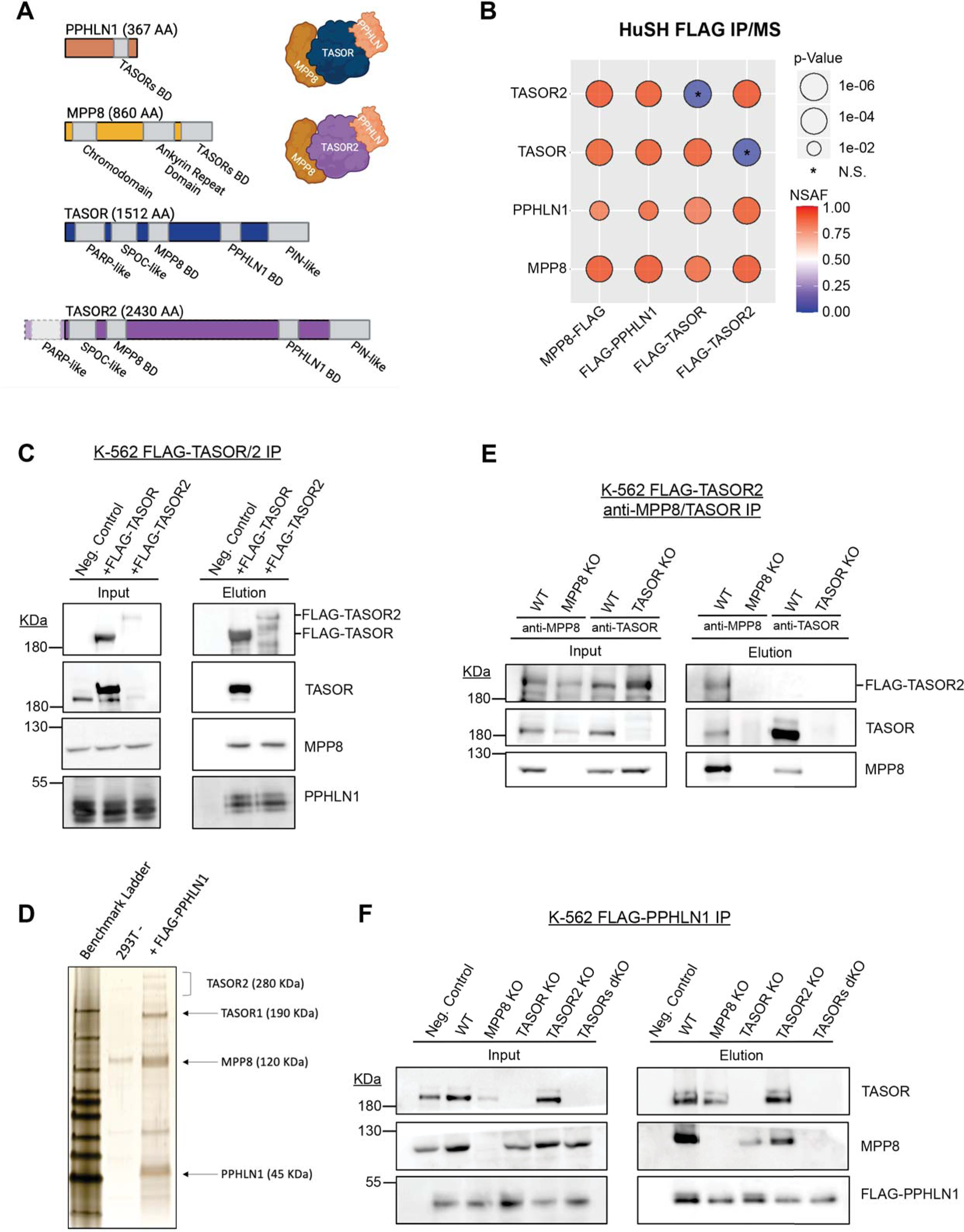
Identification of a second HuSH complex. **A)** Molecular representation of each HuSH complex member and TASOR2 and each known domain. The Binding Domain (BD) is highlighted for reference. **B)** Immunoprecipitation coupled with mass spectrometry (IP-MS) conducted in HEK293T cells expressing FLAG transgenes (x-axis) revealed the association of TASOR2 with PPHLN1 and MPP8 (y-axis). Normalized Spectral Abundance Factor (NSAF) was calculated and normalized to the length of prey targets, with comparison to the wild-type (WT) HEK293T IP negative control. IP-MS experiments were performed in bio-duplicates. **C)** Immunoblots of FLAG immunoprecipitation in K-562 cell lines expressing no transgene (WT -), FLAG-TASOR, or FLAG-TASOR2, showcasing the specific interactions in the HuSH complex. **D)** Silver stain of FLAG-PPHLN1 ammonium sulfate nuclear extract immunoprecipitation in 293F cells, providing visual evidence of the involvement of PPHLN1 in the newly discovered HuSH complex. **E)** Immunoblots demonstrating MPP8 and TASOR immunoprecipitation in parental wild-type cells, MPP8 knockouts (KOs), or TASOR KOs expressing FLAG-TASOR2, elucidating the interactions in the context of genetic perturbations. **F)** Immunoblots of FLAG immunoprecipitation performed in stable genome-edited HuSH knockout K-562 cells, expressing no transgene (Negative Control) or FLAG-PPHLN1, confirming the specificity of the interactions and the stability of the HuSH complex in a knockout background.

As expected, our analyses, which involved the use of LC-MS/MS, immunoblotting, and silver staining, validated robust associations between MPP8, PPHLN1, and TASOR. However, we made an intriguing and unexpected discovery of an association between MPP8 and PPHLN1 with an uncharacterized protein, fam208b, also known as TASOR2, which is a paralog of TASOR. By utilizing mass spectrometry analysis on samples obtained from FLAG-PPHLN1 and FLAG-MPP8 immunoprecipitation, we were able to attain a comprehensive peptide coverage across TASOR2 (**Figure S1C-D**). While TASOR and TASOR2 orthologs possess N-terminal pseudo-PARP domains, our mass spectrometry data did not reveal any peptides originating from the TASOR2 pseudo-PARP in our immunoprecipitation experiments. Notably, multiple annotated TASOR2 isoforms that lack the exons encompassing the N-terminal pseudo-PARP domain are documented. These findings indicate that TASOR2 lacking the pseudo-PARP domain is the dominant isoform within HEK293 and K-562 cells and is the isoform that associates with PPHLN and MPP8 (**Figure 1A; Figure S1A**).

To validate and further investigate the association of TASOR2 with the HuSH complex subunits, we generated stable cell lines expressing FLAG epitope-tagged TASOR and TASOR2 transgenes in K-562 cells, followed by FLAG immunoprecipitation. Our findings revealed that TASOR2 specifically associated with PPHLN1 and MPP8, while it does not co-immunoprecipitate with TASOR (**Figure 1C**). Conversely, TASOR co-IP experiments clearly demonstrated an association with PPHLN1 and MPP8, but no such association was observed with TASOR2 (**Figure 1B-C**). Additionally, an anti-MPP8 antibody efficiently co-immunoprecipitated TASOR2 in an MPP8-dependent manner. Similarly, using an anti-TASOR antibody enabled the co-immunoprecipitation of MPP8 in a TASOR-dependent manner, but no interaction was observed with TASOR2 (**Figure 1E**). These experiments collectively illustrate the existence of mutually exclusive HuSH assemblies for TASOR and TASOR2, suggesting the potential for distinct roles of these proteins within HuSH complexes.

In addition to TASOR2, our immunoprecipitation experiments revealed previously undescribed proteins that associate with HuSH assemblies. Specifically, we identified subunits of the human nuclear exosome targeting (NEXT) complex^10^, subunits of the EHMT1/2 complex^11^, and proteins involved in DNA damage response^12^ as new HuSH-associating proteins. Additionally, we observed associations between HuSH and RNA binding proteins, as well as lysine deacetylases (**Figure S1B**). Interestingly, some complexes involved in gene repression showed distinct associations with either TASOR or TASOR2. For example, the EHMT1/2 complex associated with HuSH, while subunits in lysine deacetylase complexes were associated with HuSH2. However, most associated proteins showed no distinct preference between the two HuSH complexes.

Prior work has proposed that TASOR serves as the core subunit for the HuSH complex, primarily based on the inability of MPP8 to co-IP PPHLN1 in overexpression experiments in cells^7,13,14^. Our immunoprecipitation experiments revealed that TASOR and TASOR2 exhibit mutual exclusivity in their associations with MPP8 and PPHLN1. To study the function of TASOR2 and other HuSH subunits, we employed CRISPR/Cas-9 technology to create knockout cell lines for MPP8, PPHLN1, TASOR, TASOR2, as well as a dual knockout of TASOR1/2. We found that epitope tagged PPHLN1 can associate with MPP8 in cell lines in which either TASOR or TASOR2 has been knocked out. However, the association between PPHLN1 and MPP8 was completely abrogated in the double TASOR1/2 mutant cells (**Figure 1F**). These findings indicate that TASOR and TASOR2 facilitate the association between MPP8 and PPHLN1, and that TASOR2 may serve as the core protein for a second HuSH complex. However, we observed a discernible reduction in the amount of MPP8 associating with PPHLN1 in co-IP experiments from TASOR knockout cells, when compared to TASOR2 knockout cells. This observation suggests that HuSH might be more abundant than HuSH2 within the K562 cells, also suggested by stoichiometry observed in a silver stain of both complexes (**Figure 1D**).

The results obtained from the immunoprecipitation experiments provide solid evidence for the existence of two distinct HuSH complexes, each organized around one of the two TASOR paralogs. We sought to investigate whether these two HuSH complexes bind to the same genomic loci. Prior research on HuSH has revealed its association with both pseudogenes and retrotransposable elements^2,4,8,6^. To address the question of whether the two HuSH complexes associate with identical or distinct genomic loci, we performed MPP8 chromatin immunoprecipitation sequencing (ChIP-seq) experiments. We chose the human K-562 chronic myelogenous leukemia (CML) cell line for our studies as it shows high levels of TASOR2 expression and average levels of TASOR expression relative to other commonly used cell lines (**Figure S2A-B**). We generated knockout cell lines for each HuSH subunit and conducted ChIP-seq experiments using antibodies specific for TASOR, MPP8, and PPHLN1. This allowed us to identify the genomic binding sites of the HuSH complexes and evaluate the influence of individual subunits on their localization (**Figure 2A-B, E; S3A**).

**Figure 2:**
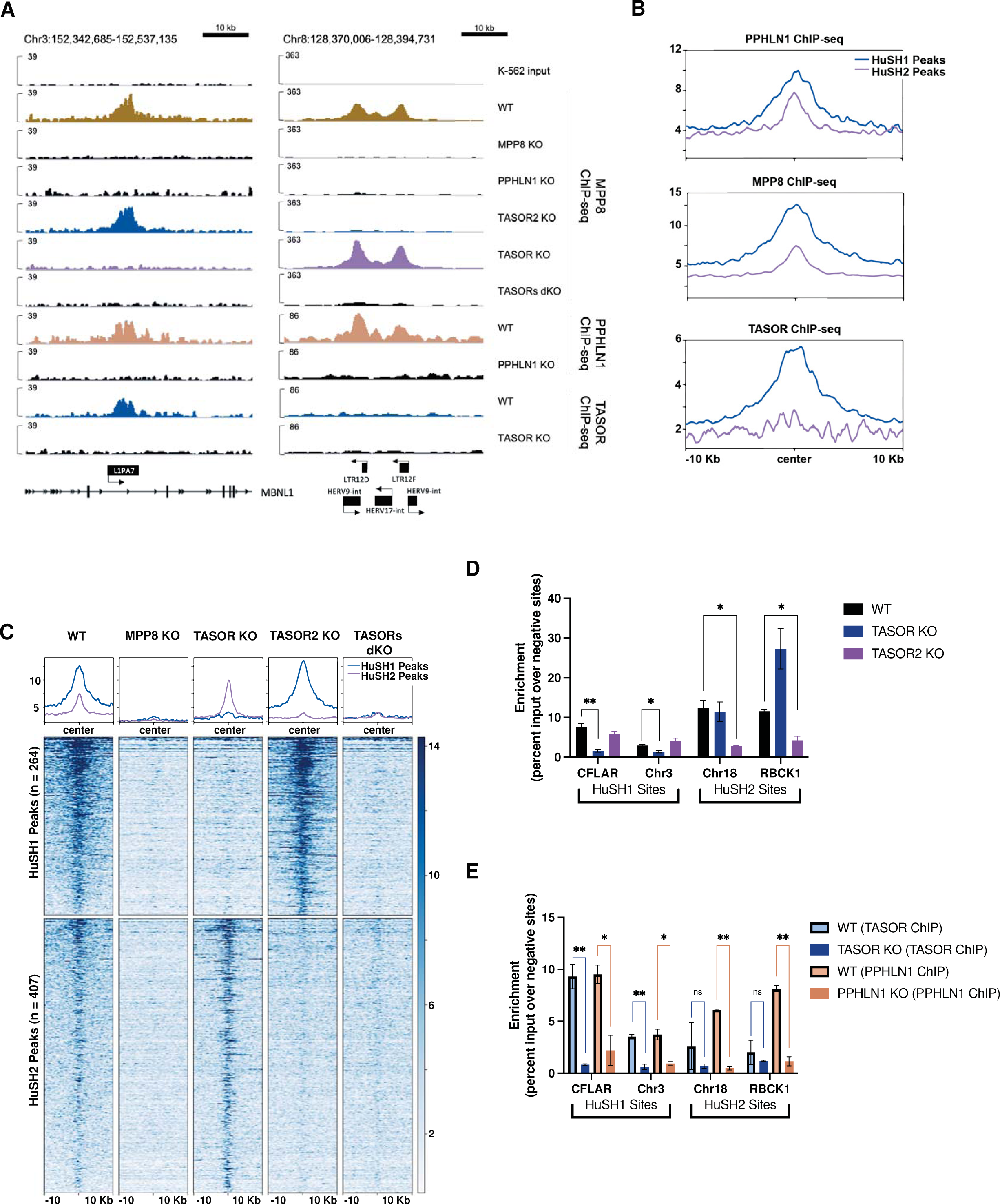
Spatial Distribution and Functional Characterization of HuSH Complexes on the Human Genome. **A)** IGV genome browser tracks depict the ChIP-seq profiles of MPP8, PPHLN1, and TASOR in both wild-type (WT) and knockout cell lines. On the left, a representative HuSH site within the L1PA7 element of MBNL1 is highlighted, while the right panel illustrates a HuSH2 site located at LTR elements. **B)** Average ChIP-seq profiles for PPHLN1, MPP8, and TASOR across HuSH and HuSH2 sites, generated using Deeptools, revealing distinct localization patterns. **C)** Heatmap and peak profiles present MPP8 ChIP-seq results in K-562 cells with knockout backgrounds. Peaks were identified using MACS2 in bio-replicates (n=2) of TASOR knockout and TASOR2 knockout, establishing two nonoverlapping regions, HuSH2 and HuSH sites, respectively. **D)** Bar chart displaying MPP8 ChIP-qPCR results with technical replicates (n=2) in knockout cell lines at HuSH and HuSH2 sites. Enrichment is measured as a percentage of input over the average percentage input at negative control sites (GNG and RABL3). Statistical analysis, compared to wild-type MPP8 peak enrichment, indicates significant (*p<0.05) or strongly significant (**p<0.01) results. **E)** Bar chart illustrating TASOR (in blue) and PPHLN1 (in peach) ChIP-qPCR outcomes in technical replicates (n=2) of wild-type and knockout cell lines at HuSH and HuSH2 sites. Enrichment is determined as a percentage of input over the average percentage input at negative control sites (GNG and RABL3). Statistical analysis, compared to knockout peak enrichment, reveals significant (*p<0.05) or strongly significant (**p<0.01) results. The data collectively highlight the distinct genomic localization and functional significance of HuSH complexes in regulating specific regions of the human genome.

Our ChIP-seq data revealed a strong correlation between the localization of MPP8 and PPHLN1 and the concurrent presence of either TASOR or TASOR2. By analyzing MPP8 and PPHLN1 ChIP-seq peaks in TASOR and TASOR2 knockout cell lines, we were able to identify specific genomic regions clearly bound by either HuSH or HuSH2 **(Figure 2C; S3A)**. Because of the unavailability of reliable TASOR2 antibodies, we have defined HuSH2 peaks as genomic regions characterized by the co-localization of both MPP8 and PPHLN1 while excluding the presence of TASOR. Additionally, these HuSH2 sites exhibit a loss of MPP8 and PPHLN1 ChIP signal in TASOR2 knockout cells, further suggesting that these sites are dependent upon the presence of TASOR2 for HuSH2 recruitment (**Figure 2A, C-D; S3A**). We observed the co-localization of all three subunits in HuSH – MPP8, TASOR, and PPHLN1 – at more than 260 genomic sites. In contrast, our analysis revealed the existence of over 400 distinct HuSH2 peaks distributed across different genomic regions. Notably, these two distinct groups exhibit absolutely no overlap in their designated genomic (**Figure S4A**). Additionally, our ChIP-seq analyses have revealed a robust correlation between HuSH and LINE1 elements, with nearly 40% of these HuSH ChIP peaks located in close proximity to LINE1 elements. This correlation is especially pronounced at the relatively young L1HS and L1PA families of primate LINE1 elements (**Figure S4B-D**). Intriguingly, this distinctive pattern of HuSH localization presents a stark juxtaposition when compared to HuSH2, which exhibit only minimal overlap with LINE1 elements. Instead of spreading throughout the genome as seen in retrotransposons such as LINE1, HuSH2 peaks are notably concentrated in close proximity to gene promoters and enhancers (**Figure S4B).**

Interestingly, we observed a significant increase in the enrichment of MPP8 and PPHLN1 at HuSH2 sites in TASOR knockout cells. Conversely, there was minimal change in the enrichment of TASOR, MPP8, and PPHLN1 at HuSH sites in TASOR2 knockout cells (**Figure 2A, C-D, S3A**). These findings substantiate the results from our co-IP experiments, suggesting higher levels of HuSH compared to HuSH2 in K-562 cells (**Figure 1D, F**). Based on the insights gained from our co-IP and ChIP-seq experiments, we propose a model in which MPP8 and/or PPHLN1 serve as critical and limiting factors, influencing the formation of the HuSH complex around either TASOR or TASOR2. To test our hypothesis regarding the competitive interaction between TASOR proteins and their influence on the localization of the two complexes, we conducted MPP8 ChIP experiments in cell lines overexpressing epitope-tagged TASOR or TASOR2 proteins (**Figure 3A-C**). Notably, the overexpression of either TASOR or TASOR2 in wildtype parental cells led to a reduction in MPP8 levels at the opposing genomic sites. These data support the proposed model, which suggests that MPP8, and potentially PPHLN1, play pivotal roles as limiting members in the assembly of functionally competent HuSH complexes.

**Figure 3:**
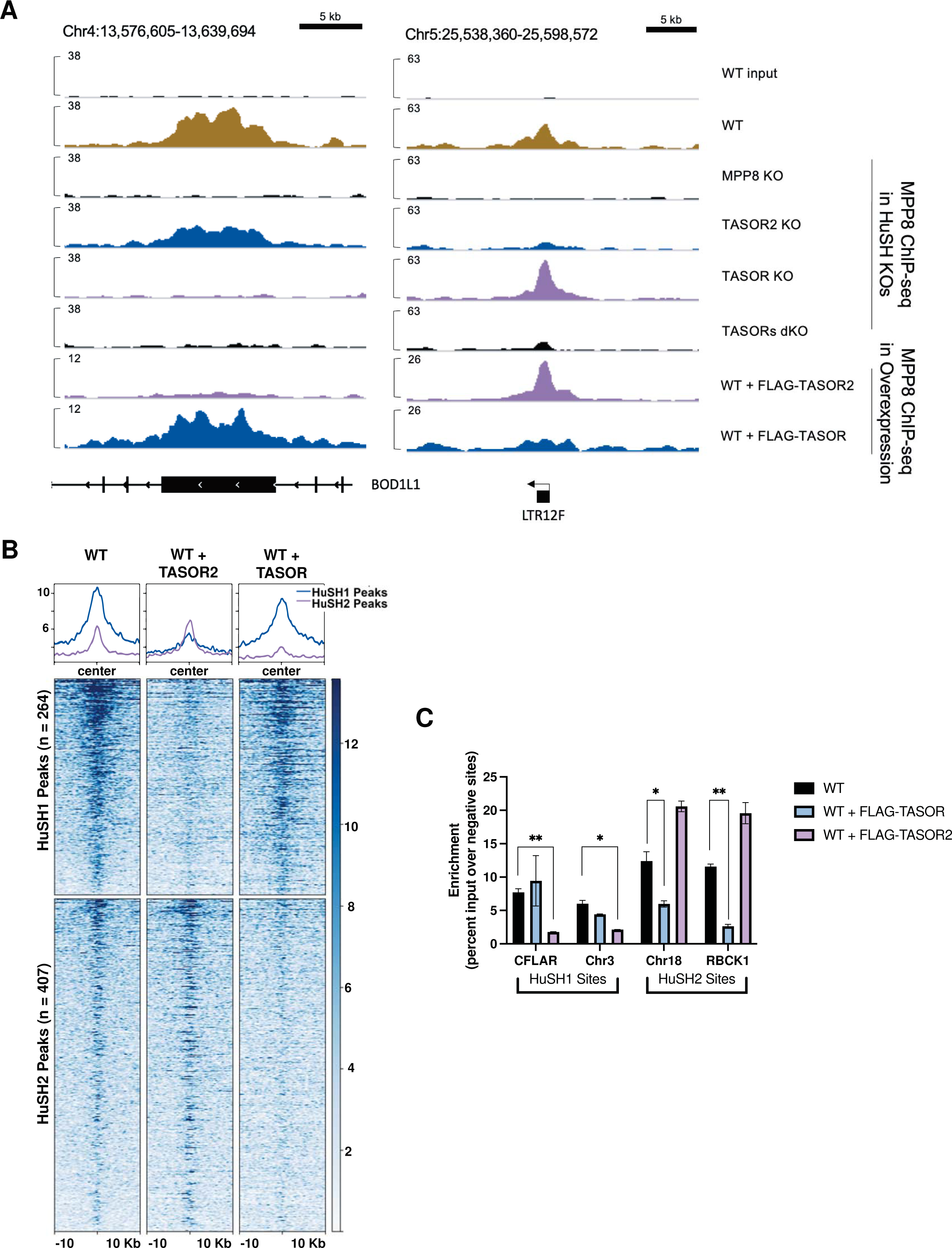
TASOR Overexpression Induces Competition for Limited MPP8 Binding. **A)** IGV genome browser tracks illustrate MPP8 ChIP-seq profiles in parental wild-type (WT) cells, knockout cell lines, and cells overexpressing FLAG-TASOR or FLAG-TASOR2. The depicted region includes one HuSH site (left) located within the exon of BOD1L1 and one HuSH2 site (right) situated at an LTR element. **B)** Heatmap and peak profiles display MPP8 ChIP-seq results from K-562 cells overexpressing FLAG-TASOR or FLAG-TASOR2. The data is sorted based on dependence on either HuSH or HuSH2. **C)** Bar chart presenting MPP8 ChIP-qPCR results from technical replicates (n=2) of FLAG-TASOR or FLAG-TASOR2 overexpressing cell lines at HuSH and HuSH2 sites. The enrichment is measured as a percentage of input over the average percentage input at negative control sites (GNG and RABL3). Statistical analysis, when compared to WT MPP8 peak enrichment, revealed significant (*p<0.05) or strongly significant (**p<0.01) results. These findings suggest a competitive relationship for MPP8 binding sites induced by TASOR overexpression.

Our ChIP-seq data provide evidence for the existence of a second HuSH complex centered around TASOR2, which exhibits distinct localization patterns within the human genome. Except for the pseudo-PARP domain, which is absent in the dominant TASOR2 isoform found in K-562 cells (**Figure S1A**), all other known domains in TASOR are present in TASOR2 (**Figure 1A**). Previous studies have suggested that the pseudo-PARP plays an important role for HuSH in LINE-1 silencing^7,15^. We aimed to explore the importance of the TASOR pseudo-PARP domain in localization of HuSH and silencing of LINE-1 in K-562 cells. We also aimed to test whether TASOR lacking its pseudo-PARP domain can relocalize to HuSH2 sites. To address these hypotheses, we introduced a wildtype TASOR transgene or a TASOR transgene lacking the pseudo-PARP domain in TASOR knockout cells and performed ChIP-seq experiments (**Figure 4A**). We performed immunoblot analysis using an antibody specific to the LINE-1 ORF1 protein (ORF1p) as a key marker for LINE-1 silencing. Our findings revealed that TASOR knockout led to the loss of TASOR ChIP-seq signal at HuSH sites, accompanied by an increase in the LINE-1 ORF1p protein (**Figure 4B-C**). Furthermore, the introduction of a wildtype TASOR transgene into TASOR knockout cells restored the ChIP-seq peaks associated with HuSH (**Figure 4C**). Remarkably, we observed that a TASOR transgene lacking the pseudo-PARP domain was unable to effectively silence LINE-1 elements in a TASOR knockout, in contrast to the wildtype transgene which demonstrated a robust decrease in ORF1p levels (**Figure 4B**). Notably, no TASOR peaks were identified in the absence of its pseudo-PARP domain, and the ChIP-seq data resembled that of a TASOR knockout. These results support the important role of the pseudo-PARP domain of TASOR in HuSH localization. However, the absence of the pseudo-PARP domain did not cause a relocalization of TASOR to HuSH2 sites. These data imply that additional domains specific to TASOR2 play an important role in the localization to gene promoters and other chromosomal regions displaying HuSH2 localization.

**Figure 4:**
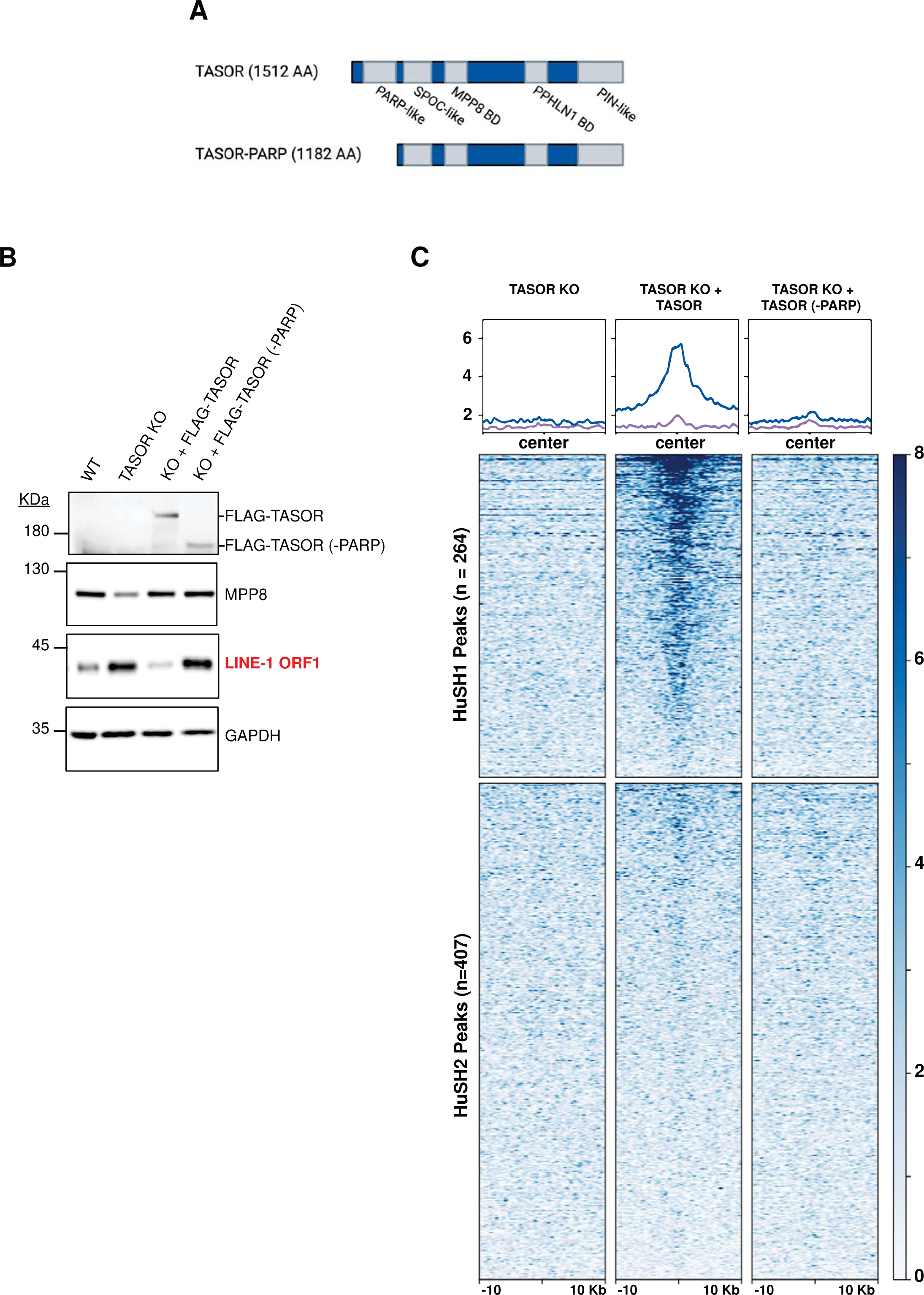
Pseudo-PARP domain is critical for HuSH localization. **A)** Molecular depiction illustrating TASOR domains and the TASOR-PARP mutant. **B)** Immunoblot analysis of whole cell lysates derived from K-562 wild-type (WT), TASOR knockout (KO), and TASOR WT or (-PARP) mutant transgene rescue cell lines. **C)** Heatmap and peak profiles illustrating TASOR ChIP-seq results conducted in TASOR KO cells with puromycin resistance, FLAG-TASOR, and FLAG-TASOR-PARP. The results are categorized into either HuSH or HuSH2 dependent peaks, shedding light on the pivotal role of the PARP domain in facilitating HuSH localization.

To investigate the roles of HuSH2 and TASOR2 in LINE-1 silencing, we evaluated ORF1p levels in HuSH subunit knockout cells (**Figure 5A**). We observed a notable upregulation of ORF1p in the PPHLN1, MPP8, and TASOR knockout cell lines, suggestive of LINE-1 silencing derepression, consistent with findings from prior research. Notably, there were no discernible changes in ORF1p expression when comparing TASOR2 knockout cells to the wildtype parental cells, consistent with the lack of enrichment of HuSH2 ChIP-seq peaks at LINE-1 elements. This finding suggests that TASOR2 may not play a direct role in the repression of LINE-1 elements within the regulatory network of the HuSH complex. In our immunoblot analysis, we detected a decrease in the steady-state levels of MPP8 and PPHLN1 in TASOR knockout cells. Conversely, the deletion of PPHLN1 or MPP8 resulted in a reduction of TASOR levels, but TASOR2 knockout cells did not exhibit a similar decrease, again suggesting lower amounts of HuSH2 in cells (**Figure 5A**).

**Figure 5:**
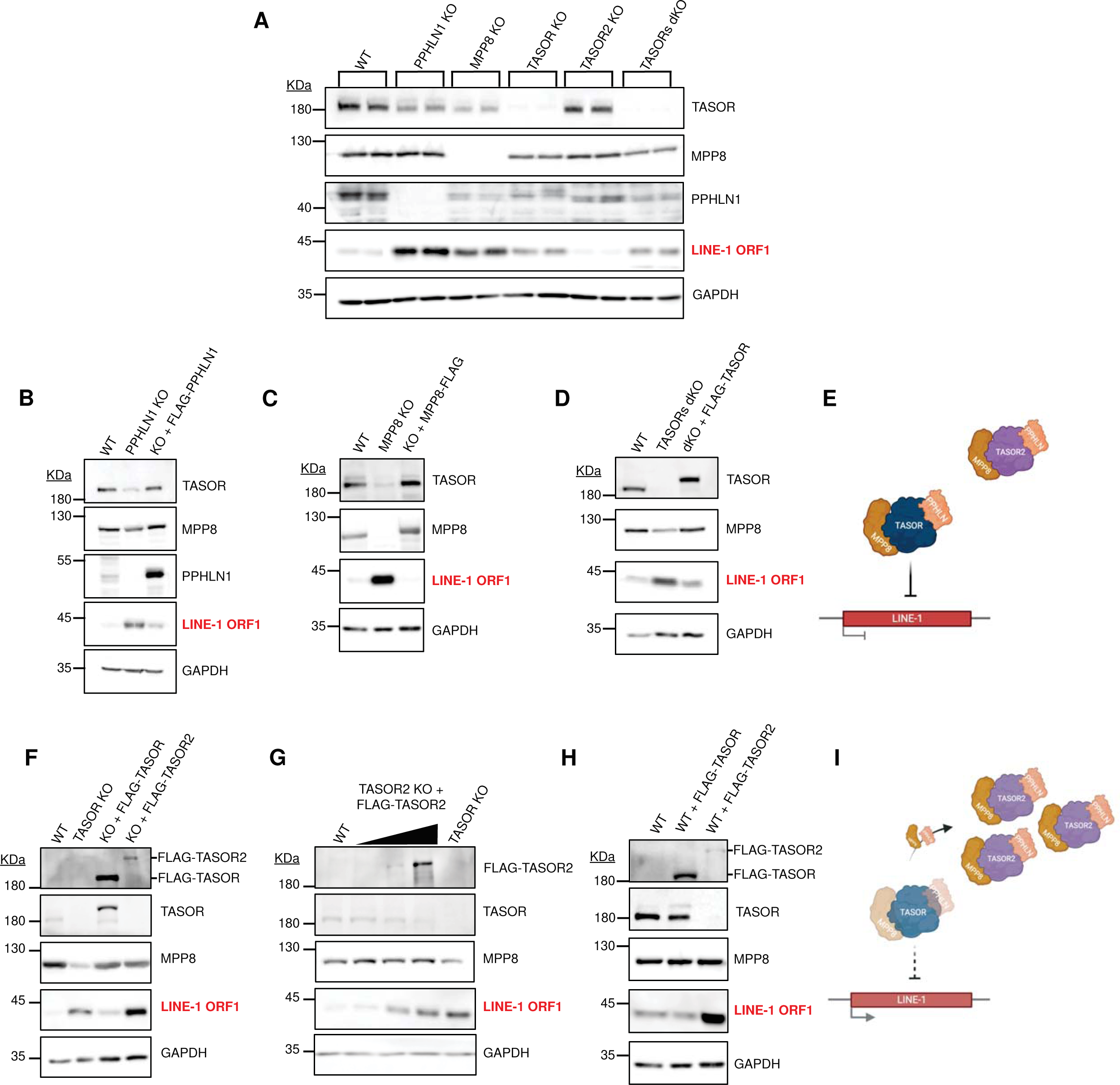
The Regulatory Dynamics of HuSH Complexes in Controlling LINE-1 Expression in K-562 Cells. **A)** Immunoblot analysis of whole cell lysates obtained from K-562 CRISPR-Cas9 stable knockouts (KOs) in bioreplicate experiments. **B-D)** Immunoblot profiles of whole cell lysates derived from K-562 wild-type (WT), knockout (KO), and FLAG transgene rescue conditions for: (B) PPHLN1 KO and FLAG-PPHLN1; (C) MPP8 KO and MPP8-FLAG; (D) TASORs double knockout (dKO) and FLAG-TASOR. **E)** Schematic representation illustrating the HuSH complex’s role in silencing LINE-1 elements. **F)** Immunoblot analysis of whole cell lysates from K-562 WT, TASOR KO, and TASOR KO with either FLAG-TASOR or FLAG-TASOR2 expression. **G)** Immunoblot analysis of whole cell lysates from K-562 WT, TASOR2 KO, with increasing levels of FLAG-TASOR2 transgene expression, compared to TASOR KO. **H)** Immunoblot analysis of whole cell lysates from K-562 WT cells overexpressing FLAG-TASOR or FLAG-TASOR2 in a wild-type background. **I)** Schematic representation depicting the dose-dependent increase of TASOR2, leading to the sequestration of MPP8 and PPHLN1 from HuSH, thereby disabling the complex from effectively silencing LINE-1 elements.

We introduced wildtype transgenes of PPHLN1, MPP8, and TASOR to investigate their potential to rescue the silencing of LINE-1 in the respective knockout models. Notably, our findings demonstrate that each of these transgenes effectively restored LINE-1 silencing in their corresponding knockout cell lines and stabilize levels of other HuSH complex members (**Figure 5B-C, F**). Furthermore, we observed that TASOR was also capable of rescuing LINE-1 silencing in a double TASOR-TASOR2 knockout cell line (**Figure 5D**). These results provide robust evidence for the regulatory role of HuSH in silencing LINE-1 elements and underscore the contributions of all subunits in this process (**Figure 5E**). In contrast, the expression of a TASOR2 transgene in a TASOR knockout background failed to rescue LINE-1 silencing but did successfully rescue MPP8 protein steady state levels (**Figure 5F**). Our co-IP and ChIP-seq findings led us to propose that PPHLN1 and/or MPP8 are limiting HuSH subunits in K-562 cells. Consequently, we suggest that competition for these two subunits could potentially regulate LINE-1 elements silencing by altering the levels of HuSH and HuSH2 in cells. An increase in HuSH2 would lead to a decrease in HuSH and loss of LINE-1 silencing (**Figure 5I**). To test this hypothesis, we titrated the levels of a wildtype TASOR2 transgene in TASOR2 knockout cells. We found that high levels of the TASOR2 transgene led to derepression of LINE-1, mimicking the effect observed in TASOR knockout cells (**Figure 5G)**. Additionally, we found that high levels of a wildtype TASOR2 transgene in wildtype K-562 cell line also led to derepression of LINE-1 elements (**Figure 5H**). Importantly, overexpression of a wildtype TASOR transgene did not have this effect. These findings reveal that by manipulating the expression levels of TASOR and TASOR2, either through knockout or overexpression, we can modulate the levels of HuSH and consequently alter the silencing of LINE-1 elements.

Our research findings indicate that MPP8 and/or PPHLN1 serve as limiting factors in the assembly of HuSH complexes. We aimed to gain a deeper understanding of how MPP8 functions in modulating the relative amounts of HuSH complexes in cells. Our hypothesis centered on the notion that disrupting the interaction between MPP8 and either TASOR or TASOR2 would result in changes in cellular levels of HuSH2 or HuSH. To achieve this, we employed AlphaFold predictions to investigate the formation of secondary and tertiary structures during the complex formation between the TASOR proteins and MPP8.

Previous studies have already defined the interaction domains between TASOR and MPP8, which include the MPP8 region located C-terminal to the ankyrin repeats, as well as the DUF3715, SPOC, and DomI domains of TASOR^7,16^. Although the DomI domains in TASOR and TASOR2 orthologs display substantial dissimilarity in their sequences, AlphaFold predictions suggest that both human TASOR proteins share a similar predicted secondary structure (**Figure 6A, Figure S5**) and adopt a structurally analogous fold that encompasses four alpha helices. One of these alpha helices is anticipated to interact with a 137-residue domain situated at the C-terminus of MPP8, comprising of beta-sheets and an alpha helix. We generated TASOR and TASOR2 transgenes with two amino acid substitutions, in which we replaced leucine or isoleucine residues with arginine. Subsequently, we introduced these transgenes into TASOR and TASOR2 knockout cells. Our results revealed that the dual L/I-to-R substitutions in both TASOR proteins disrupted their interaction with MPP8 while leaving the interaction with PPHLN1 unaffected, as demonstrated by co-immunoprecipitation (**Figure 6B, C**).

**Figure 6:**
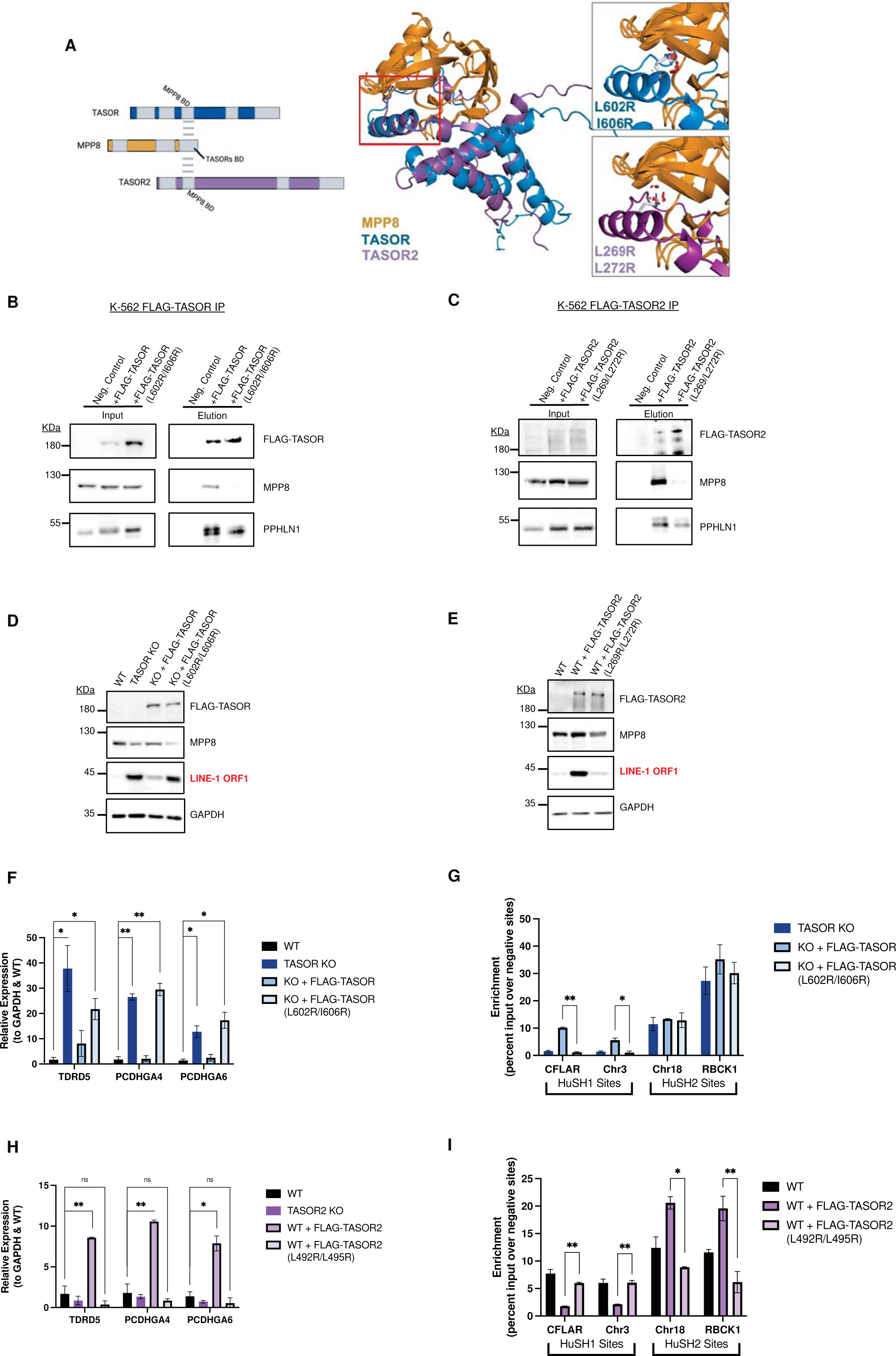
Elucidating the Binding Domain of TASOR and TASOR2 with MPP8. **A)** AlphaFold-predicted structure of the MPP8-TASORs binding domains, showcasing the overlap of DomI in both TASOR and TASOR2. Mutated residues are highlighted, signifying their predicted role in disrupting the interaction between MPP8 and TASOR or TASOR2. **B)** Immunoblots of FLAG immunoprecipitation assays conducted in TASOR knockout (KO) cell lines. These cells stably express either an empty vector (WT-), FLAG-TASOR wildtype, or a mutant transgene. **C)** Immunoblots of FLAG immunoprecipitation assays performed in TASOR2 KO cell lines, demonstrating the interaction between MPP8 and TASOR2. Cells express either an empty vector (WT-), FLAG-TASOR2 wildtype, or a mutant transgene. **D)** Immunoblot analysis of whole cell lysates obtained from K-562 wildtype (WT), TASOR KO, and WT/mutant transgene rescue cell lines, providing insights into the impact of TASOR mutations on protein expression. **E)** Immunoblot analysis of whole cell lysates from K-562 WT, TASOR2 KO, and WT/mutant transgene rescue cell lines, offering a comprehensive view of TASOR2 mutations and their effects on protein expression. **F)** Bar chart illustrating MPP8 Chromatin Immunoprecipitation followed by quantitative PCR (ChIP-qPCR) in technical replicates (n=2) of WT, TASOR KO, or WT/mutant transgene rescue cell lines. The intensity is measured at HuSH and HuSH2 sites, represented by the enrichment of percent input over the average percent input at negative control sites (GNG and RABL3). Statistical analysis, when compared to WT MPP8 peak enrichment, reveals significant (*p<0.05) or strongly significant (**p<0.01) results. **G)** Bar chart displaying MPP8 ChIP-qPCR in technical replicates (n=2) of WT or WT/mutant transgene overexpression cell lines. The intensity is measured at HuSH and HuSH2 sites by calculating the enrichment of percent input over the average percent input at negative control sites (GNG and RABL3). Statistical analysis, when compared to WT MPP8 peak enrichment, shows significant (*p<0.05) or strongly significant (**p<0.01) results, highlighting the impact of transgene overexpression on MPP8 binding.

The expression of the mutant TASOR transgene in a TASOR knockout background failed to restore LINE-1 silencing, a function effectively accomplished by expression of a wildtype TASOR transgene (**Figure 6D**). When introduced in the context of a TASOR2 transgene, the L/I-to-R substitutions impaired the ability of TASOR2 overexpression to cause derepression of LINE-1, a function attainable with overexpression of wildtype TASOR2 in parental K-562 cells (**Figure 6E**). To assess whether the derepression of LINE-1 elements is indeed linked to the inability of MPP8 to interact with and form HuSH, we performed chromatin immunoprecipitation targeting MPP8, followed by quantitative polymerase chain reaction (MPP8 ChIP-qPCR), in the TASOR mutant transgenic K-562 cell lines. TASOR knockout cells expressing the mutant TASOR transgene exhibited no MPP8 interaction at HuSH sites (**Figure 6F**). Additionally, TASOR2 knockout cells expressing a TASOR2 mutant transgene resulted in a loss of MPP8 interaction at HuSH2 sites (**Figure 6G**). These findings underscore the significance of MPP8 interaction in the functionality of the HuSH complexes.

Collectively, these findings have led us to propose a model for the silencing of LINE-1 elements by the HuSH complex. Under normal conditions in K-562 cells, TASOR acts as a central hub, facilitating the interaction between MPP8 and PPHLN1, which together form the HuSH complex. This complex associates with additional factors that may play a role in regulating its recruitment to chromatin, ultimately contributing to the subsequent silencing of LINE-1 elements. In K-562 cells, LINE-1 expression is typically low, primarily due to the high abundance of HuSH. However, when the levels of HuSH2 increase, either by reducing TASOR or increasing TASOR2 levels, MPP8 and PPHLN1 are titrated away from HuSH sites, thereby decreasing the abundance of HuSH in cells and resulting in the derepression of LINE-1 elements (**Figure 7**).

**Figure 7:**
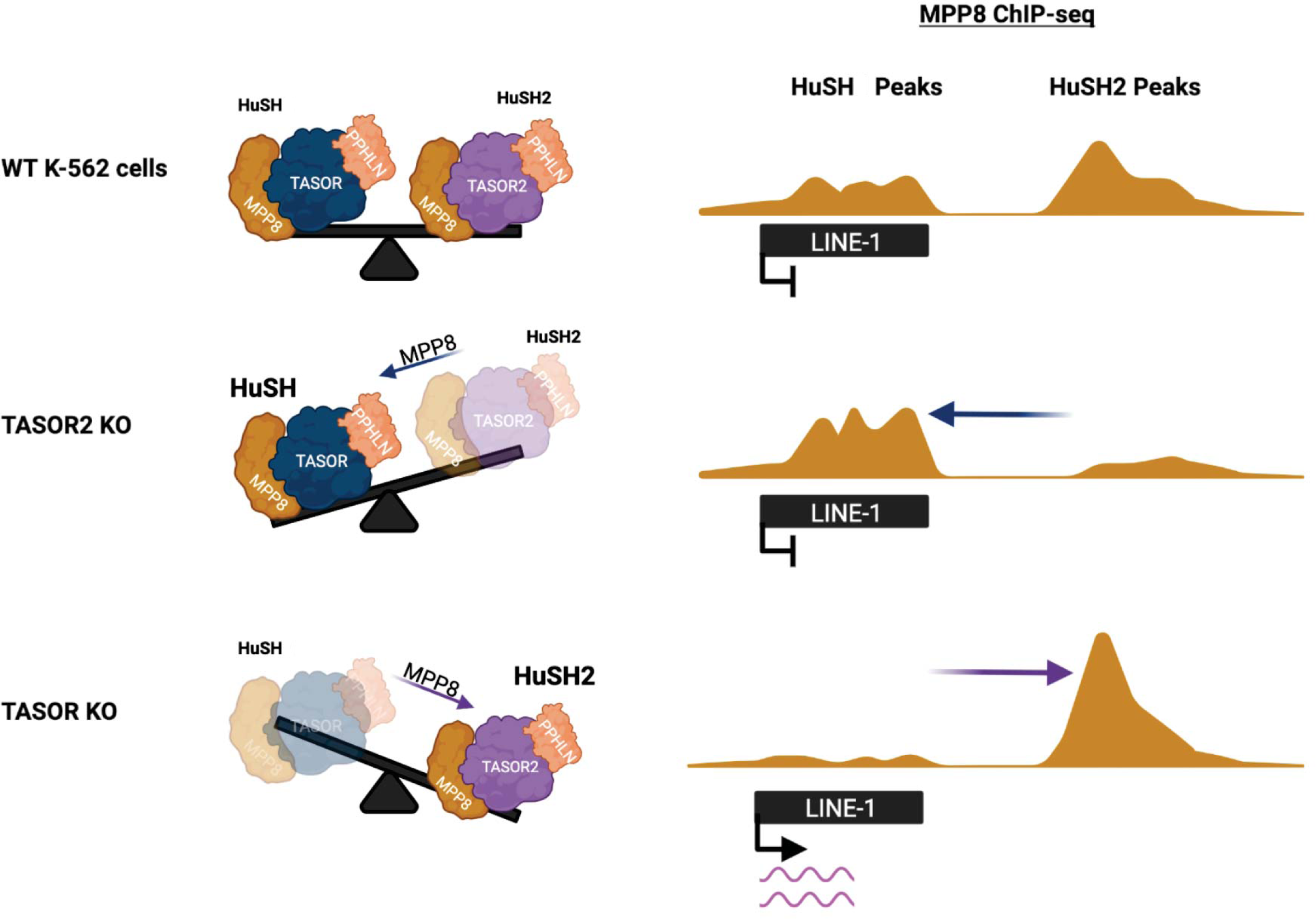
Interaction of Two Human Silencing Hub (HuSH) Complexes in Controlling LINE-1 Element Silencing. The model proposed for the silencing of LINE-1 elements by the HuSH complex is depicted in this figure. In K-562 cells under normal conditions, TASOR functions as a central hub, orchestrating the interaction between MPP8 and PPHLN1, collectively forming the HuSH complex. This complex is further associated with additional factors that likely contribute to the regulation of its recruitment to chromatin, thereby influencing the subsequent silencing of LINE-1 elements. In K-562 cells, where LINE-1 expression is typically low, the abundance of the HuSH complex plays a crucial role. However, perturbations in the balance of its components can lead to alterations in LINE-1 expression levels. Specifically, when there is an increase in HuSH2, either through the reduction of TASOR or the elevation of TASOR2, the balance of HuSH complexes begins to change. This results in the titration of MPP8 and PPHLN1, leading to a decrease in the overall abundance of HuSH. Consequently, this derepression of LINE-1 elements occurs, indicating a dynamic interplay within the HuSH complex that directly influences the regulation of LINE-1 elements in K-562 cells.

## Discussion

The identification and characterization of the HuSH complex have significantly advanced our understanding of the mechanisms orchestrating the silencing of transposable elements, particularly focusing on retrotransposable elements like LINE-1 within vertebrate animals. Our study introduces a novel variant of the HuSH complex, denoted as HuSH2, which revolves around TASOR2, a paralog of the core TASOR protein. Here, we explored the regulatory mechanisms, recruitment dynamics, and functional implications associated with both HuSH and HuSH2 complexes in the context of retrotransposon silencing.

A key revelation from our work underscores the importance of subunits within both HuSH and HuSH2 complexes for achieving genomic localization. The interconnected stability of these subunits emphasizes a meticulously regulated system, wherein alterations in TASOR and TASOR2 levels significantly influence the abundance of HuSH and HuSH2 complexes. Notably, our findings illuminate the competitive interactions between TASOR proteins for a limited pool of MPP8, emerging as pivotal factors influencing the relative quantities of HuSH complexes within cellular environments.

One of our key findings is the dynamic nature of these two complexes and the interplay between them in controlling the activity of LINE-1 elements. The changes in the abundance of HuSH and HuSH2 complexes, resulting from the competition between TASOR and TASOR2 for a limited pool of MPP8, play a pivotal role in determining the fate of retrotransposon silencing. Disturbing this delicate balance leads to the activation of transposable elements like LINE-1, underscoring the critical role played by the interplay between the two complexes.

Our ChIP-seq experiments have unveiled distinct genomic loci to which HuSH and HuSH2 complexes bind, further solidifying the existence of functionally distinct complexes. While HuSH primarily associates with LINE-1 elements and other retrotransposons, HuSH2 exhibits a distinct preference for gene promoters and enhancers. This divergence in genomic localization patterns accentuates the specialized roles these complexes play in regulating various elements within the genome. A crucial observation in our study is the pivotal role of MPP8 in governing the stability and functionality of both HuSH and HuSH2 complexes. Disrupting the interaction between MPP8 and TASOR or TASOR2 has profound implications on complex formation and, consequently, LINE-1 silencing.The presence of a pseudo-PARP domain in TASOR and TASOR2 has been of interest in previous studies, and it was suggested to be important for HuSH in LINE-1 silencing.

The presence of a pseudo-PARP domain in both TASOR and TASOR2 has been a subject of interest in previous studies, with suggestions of its importance in LINE-1 silencing by the HuSH complex. Our experiments involving TASOR lacking the pseudo-PARP domain highlight its critical role in HuSH localization and the silencing of LINE-1 elements. It is noteworthy that the absence of the pseudo-PARP domain in TASOR does not induce a relocalization to HuSH2 sites, indicating the involvement of additional TASOR2-specific domains in HuSH2 localization. Our findings also offer insights into the regulatory mechanisms governing LINE-1 silencing by HuSH and HuSH2 complexes.

In summary, our study significantly advances our understanding of the intricate interplay between HuSH and HuSH2 complexes, providing insights into their roles in the regulation of retrotransposon silencing. The competitive interactions between TASOR and TASOR2, the pivotal role of MPP8 in complex formation, and the significance of the pseudo-PARP domain in HuSH localization collectively contribute valuable knowledge to the regulatory mechanisms of HuSH complexes. These findings enrich our comprehension of how cells intricately fine-tune the silencing of retrotransposable elements, particularly LINE-1, offering a more comprehensive picture of the epigenetic regulation of the vertebrate genome.

The HuSH complex has been identified as a critical player in retrotransposon silencing across a wide range of developmental pathways and disease models. Its regulation impacts not only retrotransposons but also influences various aspects of biology. The significance of the HuSH complex in retrotransposon control has been recognized in the context of the HIV lifecycle^17–20^, as well as in mammalian development^13^. Moreover, it plays a pivotal role in the immune response in mammals^5^ and its dysregulation has been associated with Acute Myeloid Leukemia^21,22^. The intricate interplay between HuSH and HuSH2 could have significant implications for the reactivation of LINE-1 elements in development and disease. We speculate that the dynamic regulation of TASOR2 expression may be a mechanism to fine-tune and possibly attenuate HuSH-mediated silencing. This dynamic regulation of HuSH2 formation may be particularly important during certain stages of development or in specific cellular contexts, where the precise control of LINE-1 retrotransposon elements is critical. This reactivation could potentially introduce genetic instability and influence the cellular landscape. Understanding the relationship between HuSH and HuSH2 in different developmental processes and cell types may have broader implications for gene expression via regulation of retrotransposons. This hypothesis opens new possibilities for exploring the dynamic nature of gene regulation and its consequences on genome integrity.

## Acknowledgments

This work was supported by start-up funding from the University of Wisconsin School of Medicine and Public Health, the University of Wisconsin-Madison Graduate School, and the Department of Biomolecular Chemistry. Z.D.J. was supported by the National Science Foundation Graduate Research Fellowship Program under grant DGE-1747503. Any opinions, findings, and conclusions or recommendations expressed in this material are those of the authors and do not necessarily reflect the views of the National Science Foundation. P.W.L. is a Pew Scholar in the Biomedical Sciences and is supported by the National Cancer Institute (P01CA196539, R01CA266861).

## Methods

### Transgenic cell lines and culture

K-562 human lymphoblast cell lines were purchased from ATCC (CCL-243). Cells were cultured in RPMI 1640 media supplemented with 10% FBS, 1x glutamax and 1x penicillin-streptomycin, at 37°C and 5% carbon dioxide. HEK293T human embryonic kidney cells (ATCC CRL-3216) were cultured in DMEM media supplemented with 10% FBS, 1x glutamax and 1x penicillin-streptomycin, grown at 37C at 5% carbon dioxide. All cell lines were monitored for mycoplasma contamination once a month, using the LiliF e-Myco plus Mycoplasma PCR Detection Kit.

### Production of mammalian CRISPR edited cell lines

Electroporation by Amaxa nucleofector 1D machine and Lonza Nucleofector Kit V (Catalog #: VCA-1003) was used to introduce CRIPSR editing plasmid (pSpCas9(BB)-2A-GFP (PX458)) with appropriate sgRNA sequences (**Table S1**) to K-562 cells. 24 hours after electroporation cells were sorted by UW FLOW Core (Aria Cell Sorter) for single GFP cells into 96 well plates with K-562 media (described above), with an additional 10% FBS added. Successful CRISPR knockouts were screened by western blot analysis using appropriate antibodies (**Table S2**) and by genomic DNA sanger sequencing for deleterious mutations. Bio-replicates were grown separately from single cell clonal populations that had confirmed distinct mutations by use of different gRNAs or different mutations from homologous recombination confirmed through gDNA sequencing.

### Production of stable transgenic mammalian cell lines by transduction

Lentiviruses were produced by co-transfecting packaging vectors (psPAX2 and pMD2.G) and transfer vectors (pCDH-EF1a-MCS-PuroR) cloned with insertion of FLAG-PPHLN1, FLAG-TASOR, or FLAG-MPP8 transgenes, in HEK293T cells using PEI (24765-2) in vitro DNA transfection reagent. Media containing lentiviruses were collected 48 and 72 h after transfection. K-562 cells were transduced with lentiviruses for 2 days and selected using 2 ug/ml puromycin for 4 days.

### Production of stable transgenic mammalian cell lines by electroporation

For protein expression of those above lentivirus packaging limits (> 6kb DNA sequences), electroporation of pCAG vector cloned with a FLAG-TASOR2-IRES-PuroR insertion was used (Amaxa nucleofector 1D). After 24 hours K-562 cells were selected using 1 ug/ml puromycin.

### Immunoblot

Immunoprecipitation samples or whole-cell lysates were separated by SDS-PAGE, transferred to nitrocellulose membrane, blocked in 5% nonfat milk in Tris-buffered saline with 0.5 % Tween-20, probed with primary antibodies, and detected with horseradish peroxidase-conjugated or fluorescently labeled anti-rabbit or anti-mouse secondary antibodies. A list of the antibodies used is provided in **Table S1**.

### Nuclear Extract, Ammonium sulfate FLAG affinity purification

Cells were homogenized in hypotonic lysis buffer (20 mM HEPES pH 7.9, 10mM KCl, 5mM MgCl2, 0.5mM EGTA, 0.1mM EDTA 1mM DTT, 1mM Benzamidine, 0.8mM PMSF) and lysed in lysis buffer (20mM HEPES pH 7.9, 110mM KCl, 2mM MgCl2, 0.1mM EDTA, 1mM DTT, 1x Protease inhibitor cocktail, 0.4mM PMSF). Lysis was incubated in 400mM ammonium sulfate and centrifugated in ultra-centrifuge (28.5Kxg for 1.5h). Nuclear extract was dialyzed in buffer (20mM HEPES pH 7.9, 250mM KCl, 1mM EDTA, 0.01% NP-40, 0.4mM PMSF, 2mM BME) for 2 hours twice. Nuclear extract was incubated with M2 anti-FLAG affinity gel (Sigma A2220) for 2 h. Beads were washed 3-times with wash buffer (15 mM HEPES pH 7.9, 750 mM KCl, 1 mM EDTA, 0.05% Triton X-100, 8 mM PMSF) and captured proteins were eluted using 300mg/ml of 3xFLAG peptide.

### Whole cell FLAG affinity purification

Cells were lysed in buffer (20mM HEPES pH 7.9, 200mM KCl, 0.5mM EDTA, 2mM MgCl2, 1x protein inhibitor cocktail, 0.2% Triton-x, 2mM BME, 0.4mM PMSF) and the resulting lysate was incubated with M2 anti-FLAG affinity gel (Sigma A2220) for 2 h. Beads were washed 3-times with wash buffer (15 mM HEPES pH 7.9, 300-750 mM KCl, 1 mM EDTA, 0.05% Triton X-100, 8 mM PMSF) and captured proteins were eluted using 300mg/ml of 3xFLAG peptides.

### Liquid Chromatography with tandem mass spectrometry (LC-MS/MS)

After nuclear extract FLAG affinity purification, elution samples were analyzed by LC-MS/MS at the UW Biotech Center. Samples were digested in solution and then went through a desalting/clean up protocol. Normalized Spectral Abundance Factor (NSAF) calculations were utilized ^1^.

### Chromatin immunoprecipitation (ChIP)

Approximately 40 million mammalian cells were crosslinked with 1% Paraformaldehyde (Electron Microscopy Sciences) for 10 min at 37C and quenched with 0.2 M glycine. Cells were resuspended in lysis buffer (50 mM HEPES pH 7.9, 140 mL NaCl, 1 mM EDTA, 10% glycerol, 0.5% NP40, 0.25% Triton X-100). Nuclei were washed twice with digestion buffer (50 mM HEPES pH 7.9, 1 mM CaCl2, 20 mM NaCl, 1x protease inhibitor cocktail, and 0.5 mM PMSF), and treated with 105 Units of MNase (Worthington Biochemical Corporation, LS004798) per 40 million cells for 10 min at 37°C. Reaction was quenched by adding 10 mM EDTA, 5 mM EGTA, 80 mM NaCl, 0.1% sodium deoxycholate, 0.2% SDS. Nucleosomes were solubilized by sonication using Covaris S220 (160 peak incidental power, 5% duty factor, 200 cycles/burst, 45’’ ON-30’’ OFF) 6-times. One percent Triton X-100 was added to the chromatin and insoluble chromatin was removed using centrifugation. Chromatin concentration was measured using qubit and spike-in chromatin (mouse prepared in the same way) was added at a 1:40 ratio. Chromatin was incubated with primary antibodies overnight, for MPP8, TASOR and PPHLN1 (**Table S2**) 80ug of chromatin was incubated with 1.14ug antibody. Antibodies was captured using Dynabeads for 4 h at 4C and washed 3x using RIPA buffer, 2x using RIPA+300 mM NaCl and 2x with LiCl buffer. Chromatin was reverse crosslinked at 65C overnight in 10 mM Tris, 1 mM EDTA, and 1% SDS. The next day chromatin was incubated with RNase A for 1 h at 37C and proteinase K for 2 h at 55C and DNA was purified using PCR purification columns. Eluted DNA was diluted 1:10 for qPCR analysis. Sequencing libraries were prepared using NEBNext Ultra II DNA Library Prep Kit for Illumina with NEBNext Multiplex Unique Dual Oligos. Sequencing was performed on a NovaSeq6000 at the UW Biocore Sequencing Facility, paired end 150bp for 15 million reads each sample. ChIPs were performed in at least two biological replicates with similar results, data in the qPCR figures show technical replicates.

### ChIP-Sequencing analysis

Reads that passed quality score were aligned to mouse (mm9) or human (hg38) genomes using bowtie2 (parameters: -q -v 2 -m 2 -a –best –strata) (Langmead et al., 2009). Sample normalization factor was determined as ChIP-Rx = 10^6 / (total reads aligned to exogenous reference genome) or RPKM = 10^6 / (total aligned reads). Sam files were converted to bam files using samtools ^2^. Bigwig files were generated using deeptools (-bs 50 –smoothLength 600) ^3^. Peaks were called using MACS2 ^4^. Statistical analysis was performed using R. Deeptools and IGV genome browser was used for data visualization ^3,5^.

---

**Table S1.**
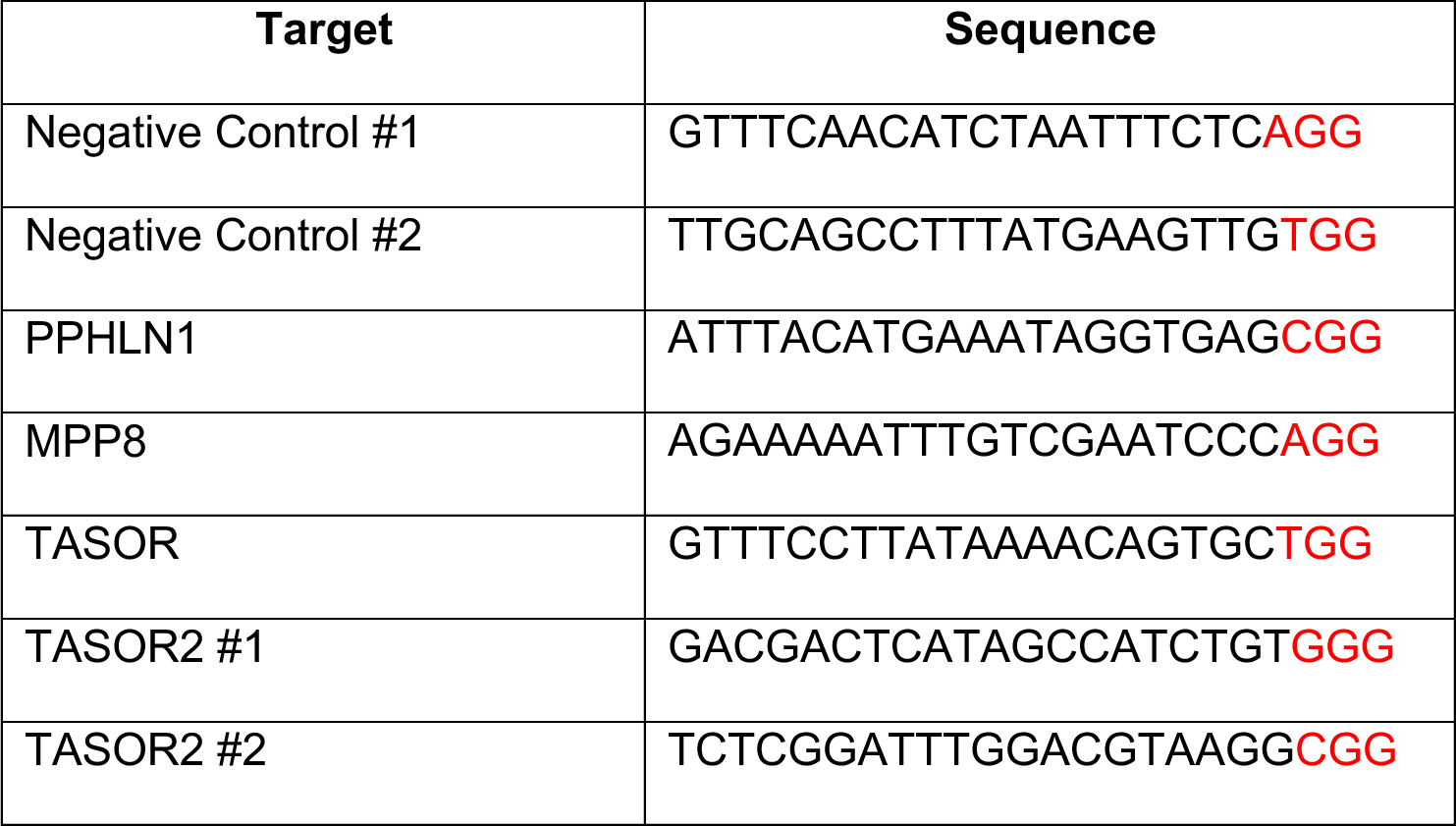
sgRNA Oligonucleotide Sequences.

**Table S2.**
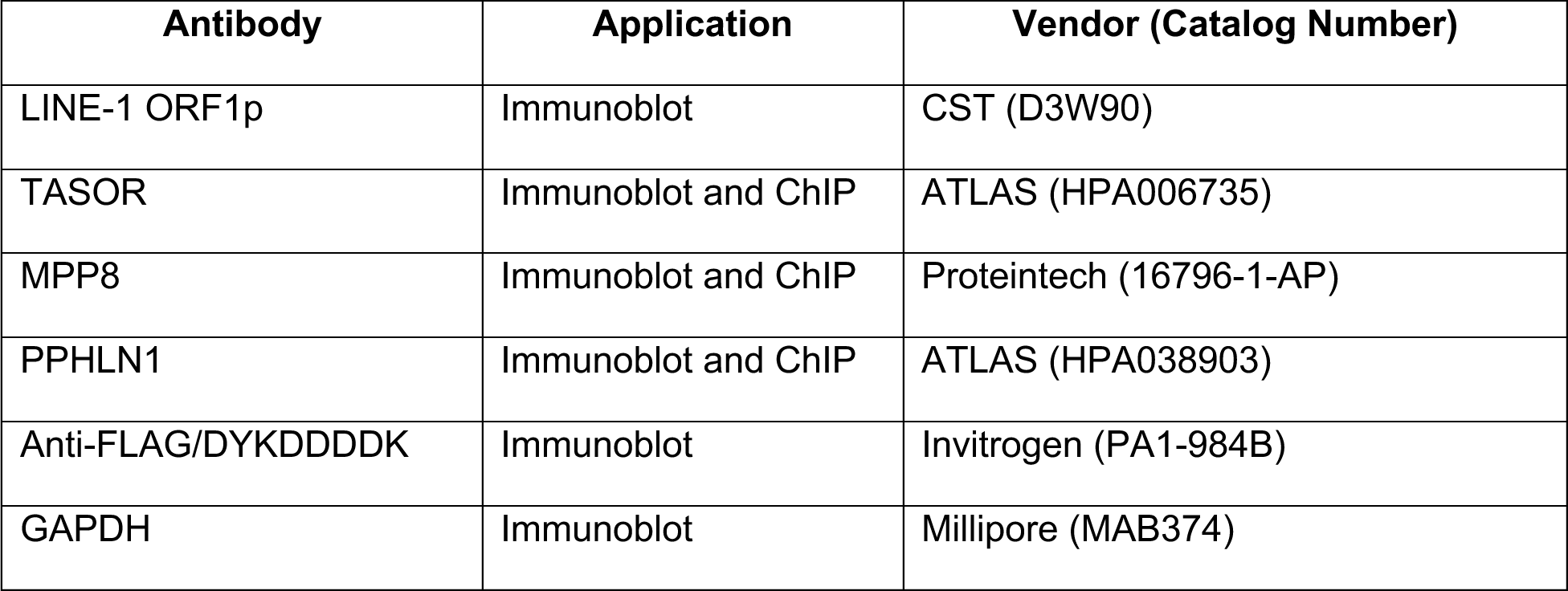
Antibodies.

| **Antibody** | **Application** | **Vendor (Catalog Number)** |
| --- | --- | --- |
| LINE-1 ORF1p | Immunoblot | CST (D3W90) |
| TASOR | Immunoblot and ChIP | ATLAS (HPA006735) |
| MPP8 | Immunoblot and ChIP | Proteintech (16796-1-AP) |
| PPHLN1 | Immunoblot and ChIP | ATLAS (HPA038903) |
| Anti-FLAG/DYKDDDDK | Immunoblot | Invitrogen (PA1-984B) |
| GAPDH | Immunoblot | Millipore (MAB374) |

**Figure S1:**
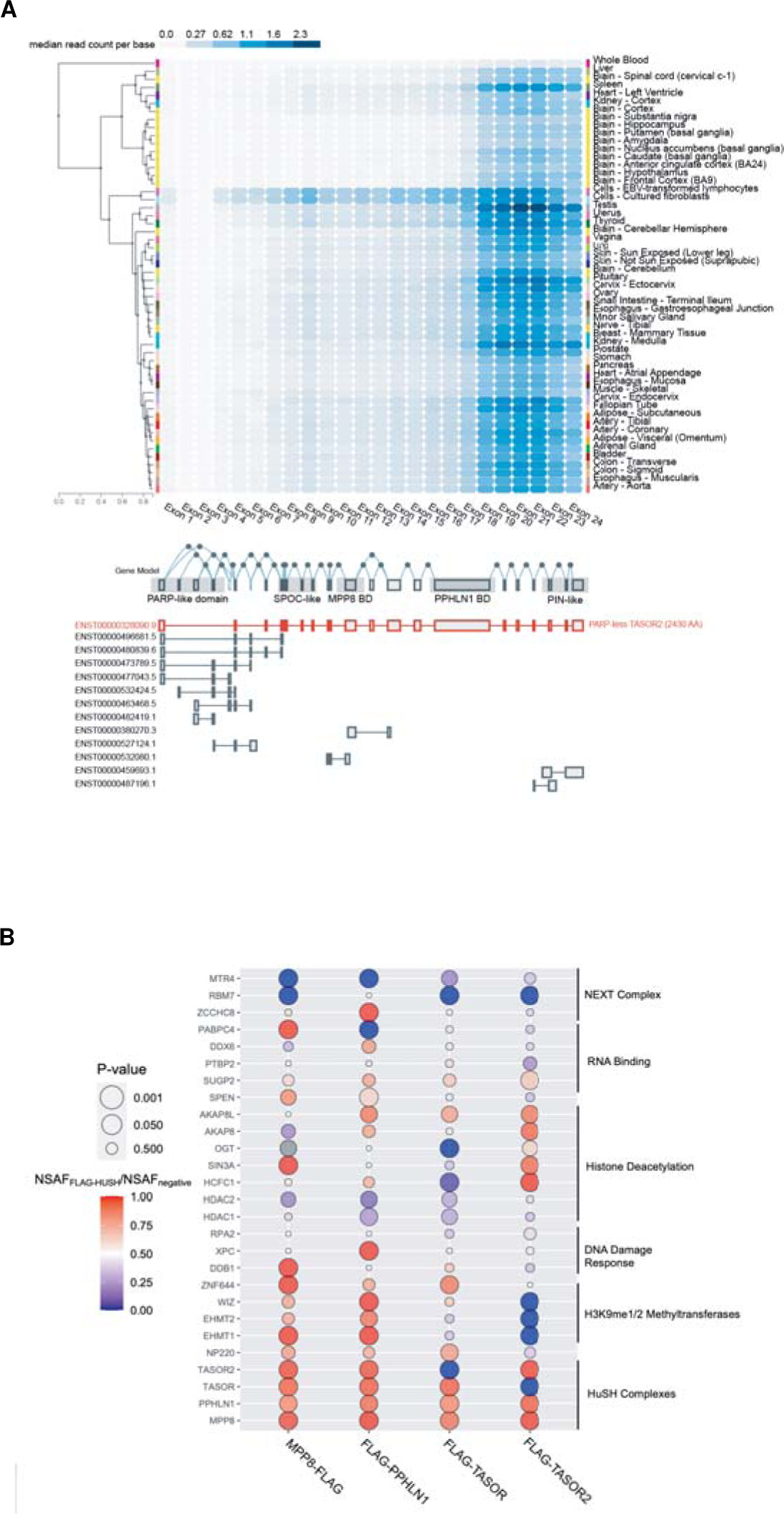
TASOR2 Isoforms and Interactions. **A)** Gene Expression Profiling in Diverse Cell Lines: Gene expression data from the Genotype-Tissue Expression (GTEx) project illustrates the differential expression of TASOR2 isoforms across various cell lines. The heatmap depicts the expression levels of distinct TASOR2 isoforms, providing insight into the tissue-specific variations. Highlighted within the figure are key domains of TASOR2, aiding in the identification and characterization of functional regions within the protein. **B)** Interactome Analysis Reveals Dynamic Complex Recruitment: Utilizing Immunoprecipitation-Mass Spectrometry/Mass Spectrometry (IP-MS/MS), the interaction landscape of TASOR2 is explored. The data reveals the recruitment of diverse protein complexes to TASOR2, particularly in association with the HuSH and HuSH2 complexes. This comprehensive analysis sheds light on the dynamic interactions that contribute to the functional versatility of TASOR2, uncovering potential regulatory mechanisms and pathways involved in its cellular roles. **C-D)** Peptide Spectra Detected by IP LC-MS/MS: Sequence of TASOR2 isoform 5 (longest isoform) including the PARP domain, with starting methionine of isoform 1 (dominant isoform expressed) without the PARP domain underlined and bolded. All peptides detected in LC-MS/MS from both FLAG-PPHLN1 and FLAG-MPP8 IPs are highlighted. C and D are two technical replicates of IP LC-MS/MS.

**Figure S2:**
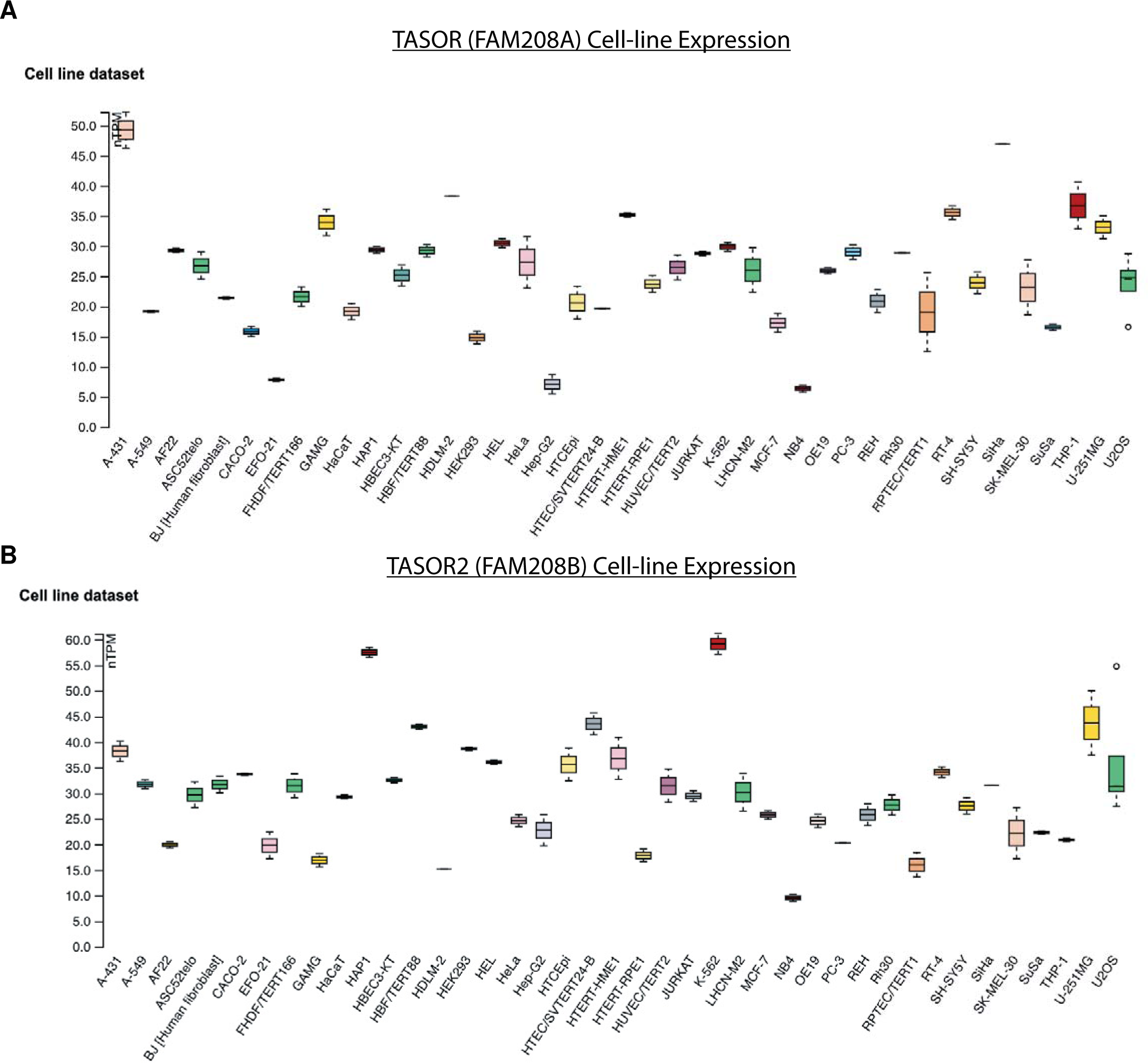
TASORs Expression in Diverse Cell Lines. **A)** The expression profile of TASOR across a spectrum of human cell lines, as determined by the Human Protein Atlas. The heatmap illustrates the varying levels of TASOR in different cellular contexts, shedding light on its dynamic presence and potential functional significance. **B)** Comprehensive analysis of TASOR2 expression in diverse cell lines, as documented by the Human Protein Atlas. The depicted data unveils the nuanced expression patterns of TASOR2 across various cellular environments, providing insights into its potential roles and regulatory mechanisms within distinct cellular contexts. The differential expression observed highlights the intricate nature of TASOR2 in cellular processes and hints at its potential implications in diverse biological pathways.

**Figure S3:**
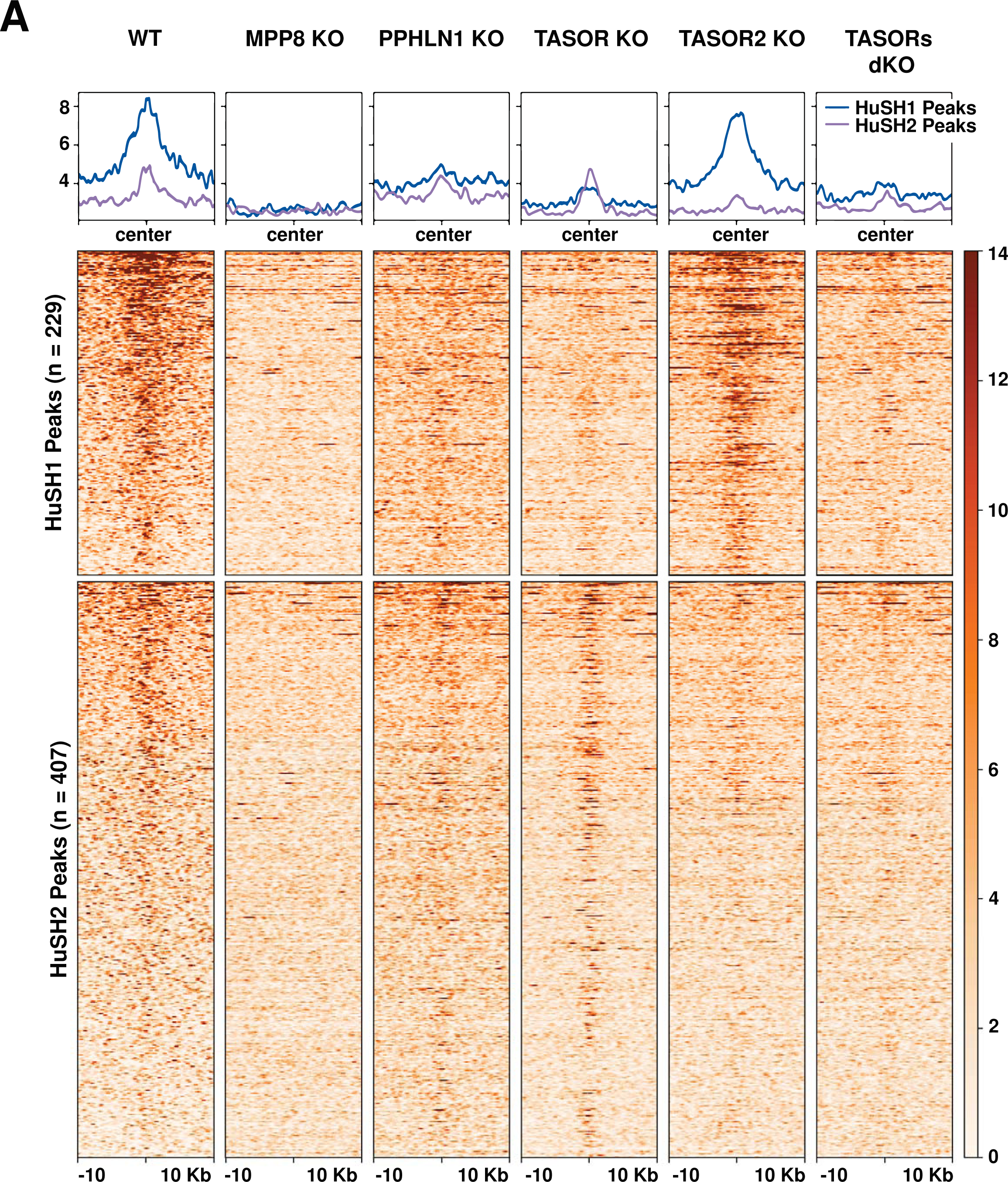
PPHLN1 ChIP-seq Analysis in HuSH KO Backgrounds. **A)** The heatmap and peak profiles illustrate the comprehensive PPHLN1 ChIP-seq results obtained from K-562 cells within the context of HuSH knockdown (KO). The ChIP-seq experiments were meticulously conducted in two distinct HuSH KO backgrounds, namely TASOR KO and TASOR2 KO, with each condition being replicated biologically (n=2). The peak calling analysis was performed using MACS2, resulting in the identification of distinct genomic regions associated with PPHLN1 binding. Specifically, two nonoverlapping sets of peaks were delineated, referred to as HuSH2 and HuSH sites, corresponding to TASOR KO and TASOR2 KO backgrounds, respectively. This segmentation provides a nuanced view of PPHLN1’s preferential binding patterns and highlights potential regulatory elements associated with the HuSH complex.

**Figure S4:**
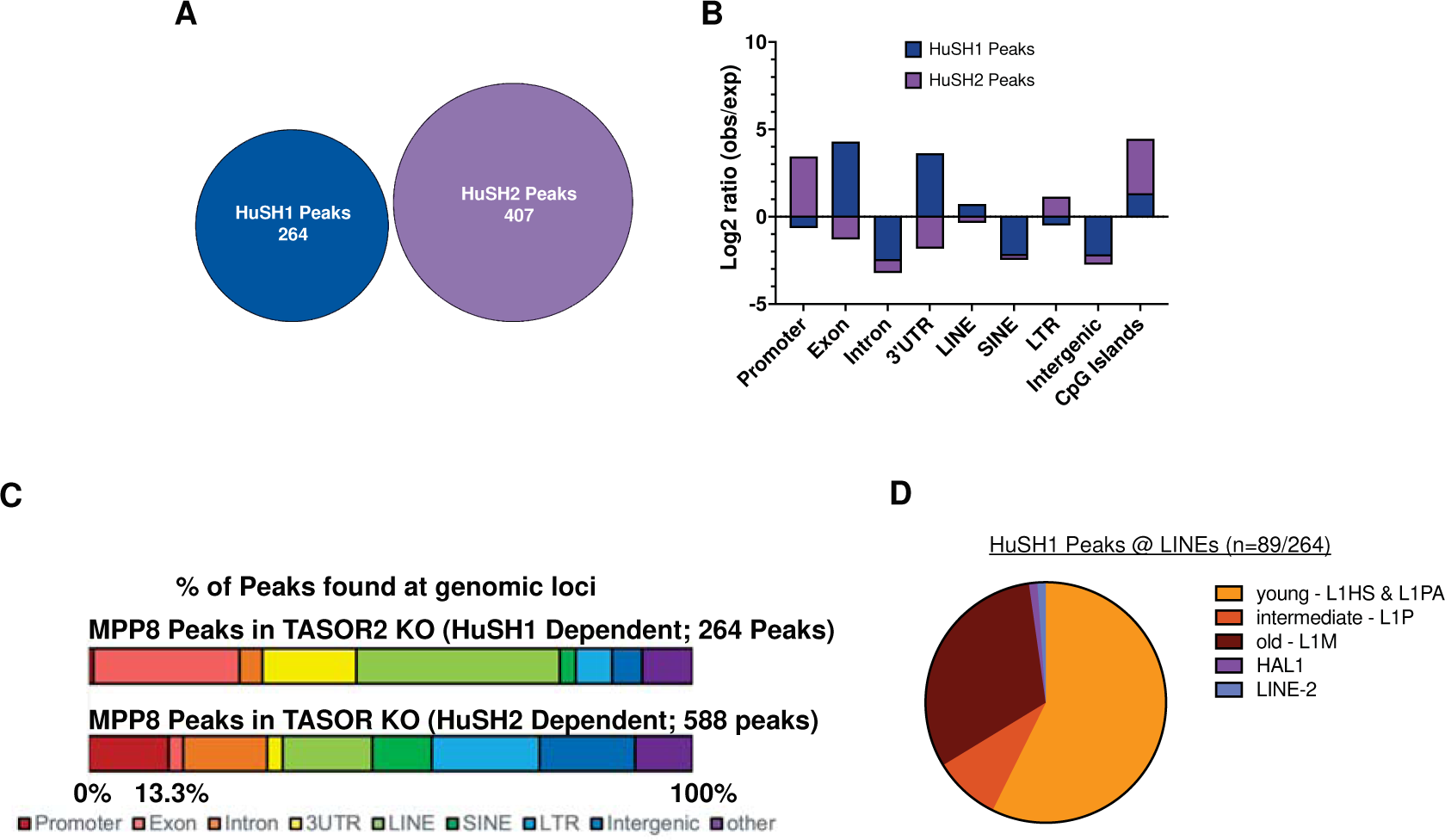
Homer Analysis Reveals Distinct Localization Patterns of HuSH and HuSH2. **A)** Venn diagram illustrating the identification of MPP8 ChIP-seq peaks designated as HuSH and HuSH2 using MACS2 in biological replicates (n=2) of TASOR KO and TASOR2 KO, resulting in the definition of two nonoverlapping regions. **B)** Bar chart presenting the observed over expected ratio of HuSH and HuSH2 peak localization at key genomic elements, as determined by Homer peak analysis. **C)** Nested bar chart depicting all MPP8 peaks found at HuSH or HuSH2 sites, categorized by their closest genomic elements. **D)** Pie chart illustrating the distribution of HuSH peaks across LINE elements of varying ages, providing insights into the preferential association of HuSH with specific LINE subtypes. This analysis contributes to a comprehensive understanding of the genomic landscape and regulatory roles of HuSH and HuSH2.

**Figure S5:**
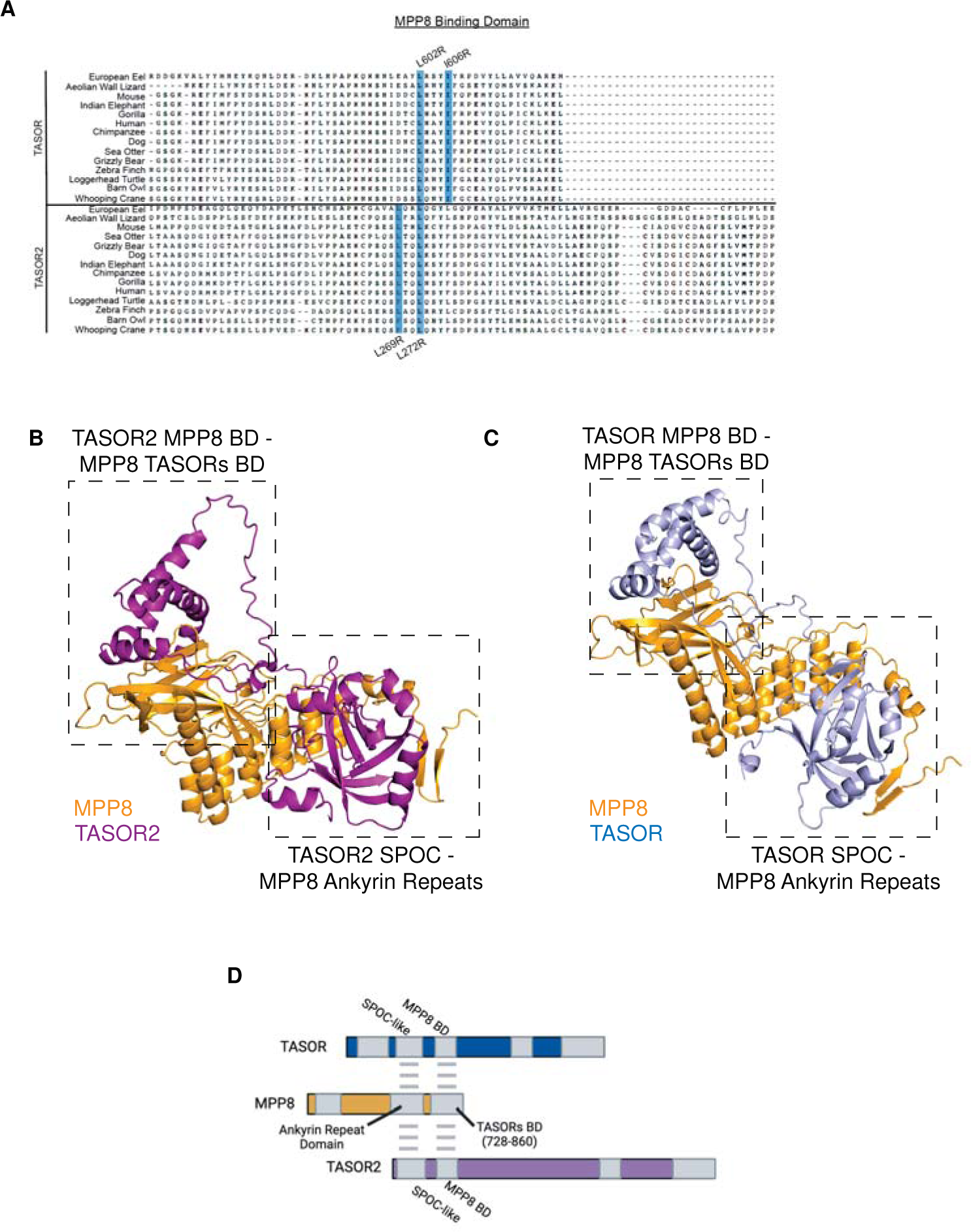
AlphaFold analysis of MPP8 interaction with TASOR/2. **A)** Protein sequence alignment of TASOR and TASOR2 orthologs depicting the conserved MPP8 binding domain. Transgenes with amino acid substitutions at the highlighted residues are illustrated in Figure 6. **B, C)** AlphaFold-predicted structure of the TASOR/2 SPOC domain interacting with the Ankyrin Repeats of MPP8. **D)** Protein domain diagram illustrating the sequences in TASOR/2 associated with MPP8.

## References

1. Tchasovnikarova, I. A. et al. Epigenetic silencing by the HUSH complex mediates position-effect variegation in human cells. Science (1979) 348, 1481–1485 (2015).

2. Tchasovnikarova, I. A. et al. Hyperactivation of HUSH complex function by Charcot–Marie–Tooth disease mutation in MORC2. Nat Genet 49, (2017).

3. Timms, R. T., Tchasovnikarova, I. A., Antrobus, R., Dougan, G. & Lehner, P. J. ATF7IP-Mediated Stabilization of the Histone Methyltransferase SETDB1 Is Essential for Heterochromatin Formation by the HUSH Complex. Cell Rep 17, 653–659 (2016).

4. Liu, N. et al. Selective silencing of euchromatic L1s revealed by genome-wide screens for L1 regulators. Nature 553, 228–232 (2018).

5. Tunbak, H. et al. The HUSH complex is a gatekeeper of type I interferon through epigenetic regulation of LINE-1s. Nat Commun 3, 54–67 (2020).

6. Seczynska, M., Bloor, S., Cuesta, S. M. & Lehner, P. J. Genome surveillance by HUSH-mediated silencing of intronless mobile elements. Nature 2021 601:7893 601, 440–445 (2021).

7. Douse, C. H. et al. TASOR is a pseudo-PARP that directs HUSH complex assembly and epigenetic transposon control. Nature Communications 2020 11:1 11, 1–16 (2020).

8. Robbez-Masson, L. et al. The HUSH complex cooperates with TRIM28 to repress young retrotransposons and new genes. Genome Res 28, 836–845 (2018).

9. Spencley, A. L. et al. Co-transcriptional genome surveillance by HUSH is coupled to termination machinery. Mol Cell 83, 1623–1639.e8 (2023).

10. Garland, W. et al. Chromatin modifier HUSH co-operates with RNA decay factor NEXT to restrict transposable element expression. Mol Cell 82, 1691–1707.e8 (2022).

11. Tsusaka, T. et al. Tri-methylation of ATF7IP by G9a/GLP recruits the chromodomain protein MPP8. Epigenetics Chromatin 11, 56 (2018).

12. Marnef, A. et al. A cohesin/HUSH-and LINC-dependent pathway controls ribosomal DNA double-strand break repair. Genes Dev 33, 1–16 (2019).

13. Müller, I. et al. MPP8 is essential for sustaining self-renewal of ground-state pluripotent stem cells. Nat Commun 12, (2021).

14. Prigozhin, D. M. et al. Periphilin self-association underpins epigenetic silencing by the HUSH complex. Nucleic Acids Res 48, 10313–10328 (2020).

15. Schöpp, T. et al. The DUF3715 domain has a conserved role in RNA-directed transposon silencing. RNA rna.079693.123 (2023) doi:10.1261/RNA.079693.123.

16. Müller, I. et al. MPP8 is essential for sustaining self-renewal of ground-state pluripotent stem cells. Nat Commun 12, (2021).

17. Matkovic, R. et al. TASOR epigenetic repressor cooperates with a CNOT1 RNA degradation pathway to repress HIV. RNA 29, 1471–1480 (2023).

18. Chougui, G. et al. HIV-2/SIV viral protein X counteracts HUSH repressor complex. Nat Microbiol 3, 891– 897 (2018).

19. Machida, S. et al. Exploring histone loading on HIV DNA reveals a dynamic nucleosome positioning between unintegrated and integrated viral genome. Proc Natl Acad Sci U S A 1–9 (2020) doi:10.1073/pnas.1913754117.

20. Yurkovetskiy, L. et al. Primate immunodeficiency virus proteins Vpx and Vpr counteract transcriptional repression of proviruses by the HUSH complex. Nat Microbiol 3, 1354–1361 (2018).

21. Cuellar, T. L. et al. Silencing of retrotransposons by SETDB1 inhibits the interferon response in acute myeloid leukemia. J Cell Biol 216, 3535–3549 (2017).

22. Gu, Z. et al. Silencing of LINE-1 retrotransposons is a selective dependency of myeloid leukemia. Nat Genet (2021) doi:10.1038/s41588-021-00829-8.

## References

1. Zhang, Y., Wen, Z., Washburn, M. P. & Florens, L. Refinements to label free proteome quantitation: How to deal with peptides shared by multiple proteins. Analytical Chemistry 82, 2272–2281 (2010).

2. Danecek, P. et al. Twelve years of SAMtools and BCFtools. GigaScience 10, 1–4 (2021).

3. Ramírez, F. et al. deepTools2: a next generation web server for deep-sequencing data analysis. Nucleic Acids Research 44, W160–W165 (2016).

4. Zhang, Y. et al. Model-based analysis of ChIP-Seq (MACS). Genome Biology 9, 1–9 (2008).

5. Robinson, J. T. et al. Integrative Genomics Viewer. Nature biotechnology 29, 24 (2011).

